# Liver x receptor alpha drives chemoresistance in response to side-chain hydroxycholesterols in triple negative breast cancer

**DOI:** 10.1101/2020.08.04.235697

**Authors:** Samantha A Hutchinson, Alex Websdale, Giorgia Cioccoloni, Hanne Røberg-Larsen, Priscilia Lianto, Baek Kim, Ailsa Rose, Chrysa Soteriou, Laura M Wastall, Bethany J Williams, Madeline A Henn, Joy J Chen, Liqian Ma, J Bernadette Moore, Erik Nelson, Thomas A Hughes, James L Thorne

**Author notes:** corresponding authors Dr James L Thorne; School of Food Science and Nutrition, University of Leeds, Leeds, LS2 9JT, UK; 0044 113 3430684; and, Dr Thomas A Hughes; School of Medicine, University of Leeds, Leeds, LS9 7TF, UK; 0044 113 3431984. these authors contributed equally to the work in the manuscript (Samantha A Hutchinson, Alex Websdale).

## Abstract

Triple negative breast cancer (TNBC) is challenging to treat successfully because targeted therapies do not exist. Instead, systemic therapy is typically restricted to cytotoxic chemotherapy, which fails more often in patients with elevated circulating cholesterol. Liver x receptors are ligand-dependent transcription factors that are homeostatic regulators of cholesterol, and are linked to regulation of broad-affinity xenobiotic transporter activity in non-tumor tissues. We show that LXR ligands confer chemotherapy resistance in TNBC cell lines and xenografts, and that LXRalpha is necessary and sufficient to mediate this resistance. Furthermore, in TNBC patients who had cancer recurrences, LXRalpha and ligands were independent markers of poor prognosis and correlated with P-glycoprotein expression. However, in patients who survived their disease, LXRalpha signaling and P-glycoprotein were decoupled. These data reveal a novel chemotherapy resistance mechanism in this poor prognosis subtype of breast cancer. We conclude that systemic chemotherapy failure in some TNBC patients is caused by co-opting the LXRalpha:P-glycoprotein axis, a pathway highly targetable by therapies that are already used for prevention and treatment of other diseases.

## Introduction

Breast cancer (BCa) prognosis depends on tumour and host parameters. Breast tumors that don’t express the estrogen (ER), progesterone (PR), and Her2 receptors (ER-/PR-/Her2-), termed triple negative (TNBC), are challenging to treat successfully because therapies such as Tamoxifen and Herceptin that target ER and Her2 signaling respectively, are not effective. Typically, TNBC patients undergo surgical resection of primary tumor and lymph nodes, radiotherapy, and systemic cytotoxic chemotherapy. Chemotherapy can be given either before (neoadjuvant [NACT]) or after (adjuvant [ACT]) surgery.

NACT is increasingly common to downstage the tumor to allow breast conserving surgery to be performed (Bagegni *et al*, 2019). A clinically measurable surrogate marker of likely outcome following NACT is pathological complete response (pCR) to treatment (Xia *et al*, 2020). If pCR is observed after NACT then prognosis is generally very good and invasive surgery is minimized. If pCR is not observed then treatment and can be adapted, but prognosis is worse and patients are likely to have been better placed on alternative treatments, such as long term adjuvant chemotherapy after surgical resection (Xia *et al*., 2020). If TNBC patients are going to relapse with their disease, they typically do so within 2-3 years (Liedtke *et al*, 2008), highlighting a substantial subset of patients in whom systemic therapy has failed and who now have few remaining treatment options. Maximizing response to chemotherapy is a critical clinical need, yet the mechanisms responsible for resistance remain poorly understood (Eccles *et al*, 2013).

Circulating cholesterol levels appear to be inversely linked to survival of BCa patients, and are extensively modifiable by diet, lifestyle, and pharmacological interventions. Survival is worse for BCa patients who present with high LDL-cholesterol (dos Santos *et al*, 2014), but is improved with dietary (Brennan *et al*, 2017; Chlebowski *et al*, 2006; Jiang *et al*, 2019) or pharmaceutical (Liu *et al*, 2017b) regimens that are associated with normalized LDL-C. A clinical trial that promoted dietary methods to reduce LDL-cholesterol was found to reduce breast cancer recurrence rates (Toledo *et al*, 2015). Robust mechanistic evidence explaining these links remains sparse, thus the guidance that can be offered to different patient groups with respect to the potential benefits of reducing LDL-cholesterol remains limited (De Cicco *et al*, 2019). Cholesterol is the precursor for an array of compounds including hormones, seco-steroids, and bile acids. During the synthesis of these compounds, a diverse array of oxysterol intermediates are produced that are potent signaling molecules in their own right. Hydroxylation of the cholesterol side chain produces side-chain hydroxycholesterols (scOHC), such as 24-hydroxycholesterol (24OHC), 26-hydroxycholesterol (26OHC/27OHC), epoxycholesterols (e.g. 24,25-epoxycholesterol), and oxysterol conjugation can produce a wider variety of signaling molecules such as dendrogenin A. Oxysterols are continually detected by the liver x receptor alpha (LXRα) and liver x receptor beta (LXRβ) transcription factors (Janowski *et al*, 1996), allowing for homeostatic control of end product synthesis and simultaneously inhibiting potentially harmful build-up of oxysterol intermediates (Clare *et al*, 1995) or of cholesterol itself (Tabas, 2002). Selective modulation of LXRα and LXRβ by oxysterols leads to divergent effects in cancer pathophysiology. For example, the oxysterol-histamine conjugate dendrogenin A preferentially activates LXRβ, induces lethal autophagy (Poirot & Silvente-Poirot, 2018) and differentiation of breast cancer cells (Bauriaud-Mallet *et al*, 2019).

Contrary to this, in TNBC scOHCs promote metastatic colonization (Baek *et al*, 2017) and epithelial mesenchymal transition, and in ER-positive breast cancers, scOHCs both reduce endocrine therapy efficacy (Nguyen *et al*, 2015) and are circulating biomarkers of recurrence (Dalenc *et al*, 2017).

Interestingly, there is evidence that LXRα and scOHCs may also play a role in chemotherapy resistance. LXRα maintains integrity of the blood brain barrier (Wouters *et al*, 2019) via upregulation of the multi-drug resistance pump, P-glycoprotein, in response to oxysterols (Saint-Pol *et al*, 2013) and synthetic LXR ligands (ElAli & Hermann, 2012). The ATP-Binding Cassette B1 (ABCB1)/P-glycoprotein (Pgp) is a highly promiscuous xenobiotic efflux transporter with substrate diversity that includes cholesterol (Garrigues *et al*, 2002) and an array of chemotherapy agents typically given to TNBC patients. LXR and Pgp therefore potentially link oxysterol signaling to chemotherapy efficacy. We previously demonstrated that ER-negative breast cancer was highly responsive to LXRα signaling compared to other breast cancer types (Hutchinson *et al*, 2019) suggesting retention of the scOHC:LXR axis provided a survival advantage to TNBC tumors that persisted despite the anti-proliferative actions LXR confers. Here, we have explored the hypothesis that failure of systemic chemotherapy could be attributed to a cholesterol rich tumour environment, where aberrant activation of the scOHC:LXR:Pgp signalling axis is co-opted to confer resistance to common chemotherapy drugs.

## Materials and Methods

### Cell culture and cell lines

Cell lines were originally obtained from ATCC. All cells were routinely maintained at 37°C with 5% CO2 in Dulbecco’s Modified Eagle Medium (DMEM, Thermo Fisher, Cat: 31966047) supplemented with 10% FCS (Thermo Fisher, UK, Cat: 11560636). Cell lines were confirmed mycoplasma free at 6-monthly intervals, and cell line identities were authenticated at the start of the project.

### Drugs and reagents

All stocks were stored at -20°C. scOHCs were from Avanti (Alabama, US) and stored as 10 mM stocks in nitrogen flushed ethanol: 24OHC (#700071), 25OHC (#700019), 26OHC (#700021). Epirubicin (Cayman, UK Cat:12091) was stored at 10 mM in nuclease free water and protected from light. GSK2033 (ToCris, Abindon, UK – #5694) at 20 mM diluted in ETOH. ABC inhibitors: MK-571 (Cambridge Bioscience – Cat: 10029-1mg-CAY) and KO143 (Sigma – Cat: K2144-1mg) were diluted in DMSO, while Verapamil (Insight Biotechnology – Cat: sc-3590) was diluted in NFW; all at 10 mM. TaqMan assays (Thermo Fisher, Paisley, #4331182): LXRα [Hs00172885_m1] LXRβ [Hs01027215_g1], Pgp [Hs00184500_m1], ABCA1 [Hs01059137_m1], HPRT1 [Hs02800695_m1]. Origine trisilencer complexes (Maryland, US): LXRα #SR322981, LXRβ #SR305039). Antibodies: Pgp (Santa Cruz Biotech, CA, US - #sc73354), CYP27A1 (Abcam, Cambridge, UK - #ab126785), CYP46A1 (Abcam, Cambridge, UK - #ab198889), CH25H (Bioss, MA, US - #bs6480R), LXRα (R&D System, Minneapolis, US - #PP-PPZ0412-00), LXRβ (Active Motif, Carslbad, US - #61177). Antibody validation is described in detail in Supplementary Information.

### Analysis of gene expression

Analysis of gene expression was performed as described previously (Hutchinson *et al*., 2019). Briefly, mRNA was extracted using Reliaprep Minipreps for cell cultures (Promega, UK, #Z6012) and RNA Tissue Miniprep System (Promega, UK, #Z6112 for tumor samples. GoScript^™^ (Promega, UK, #A5003) was used for the cDNA synthesis. Taqman Fast Advanced Mastermix (Thermo Fisher, Paisley, #4444557) was mixed with Taqman assays and analysed in a 384 well QuantStudio Flex 7 (Applied Biosystems Life Tech, Thermo Scientific) in 5μl reaction volumes. siRNA (30nM) were transfected with RNAiMAX (Thermo Fisher, #13778030) according to manufacturer’s instructions with following adaption. Cells were incubated for 22 h with reagent/siRNA mixture and then media was exchanged. After a further 14 h RNA was extracted. All experiments were conducted in technical triplicate and are presented as mean ± SEM of three or more independent replicates

### Cell viability assays

MTT were performed as previously described (Hutchinson *et al*., 2019). Briefly, cells were suspended to 1×10^6^ cells/mL, 250 μL/well of cell suspension was plated into 6-well plates and topped up to 2 mL with DMEM-10%. Cells were incubated overnight and then treated with GW3965, GSK2033, scOHC or vehicle control (VC) (ETOH/DMSO/N^2^F ETOH) for 24 h. Epirubicin (25 nM) or VC (nuclease free water) was added for a further 24 h. Cells were then counted and 500 cells per treatment were plated in triplicate in 6 well plates (Nunc, Thermo Fisher, UK, Cat: 10119831) and incubated for 12 days. Colonies were washed with PBS and stained with a 0.1 % crystal violet staining solution was this in 50 % methanol, 30 % ethanol and 20 % ddH_2_0 (Sigma, UK, Cat: V5265-250 ML). Colonies were left to air dry overnight then counted. All experiments were conducted in technical triplicate and are presented as mean ± SEM of three or more independent replicates

Colony forming assays (CFA) were performed as previously described (Thorne *et al*, 2018). Briefly, 2 ⨯10^4^ cells were seeded in 96 well plates and incubated overnight. For the combination treatment with EPI, cells were pre-treated with LXR synthetic ligands (0.1, 0.25, or 1 μM), or scOHC (1, 2.5, or 10 μM) for 24 h. After 24 h treatment, EPI was added and incubated for the next 48 h. Cells were washed with 100 μL PBS/well. 90 μL phenol red free DMEM supplemented with 10% FBS was added to each well and followed with the addition of 10 μL of diluted MTT reagent (Sigma-Aldrich, UK, Cat: M2128; final concentration 0.5 mg/mL) to each well. After 4 h incubation at 37°C, media was carefully removed and replaced with 100 μL of DMSO/well. The absorbance was read using a CLARIOstar plate reader (BMG LABTECH, Germany) at 540 nm. All experiments were conducted with six technical replicates and are presented as mean ± SEM of three or more independent replicates.

### Chemotherapy efflux Assay

Cells were plated (50,000 cells/well) in clear bottom black walled tissue culture 96 well plates (Greiner Bio-One, UK, Cat: 655986) and pre-treated with either VC (ETOH) or LXR ligands (GSK2033, GW3965 at 1 µM, 24OHC, 26OHC, 10 µM) for 16 h before a high dose of chemotherapy agent (50 µM epirubicin) for 1 h. Cells were gently washed with PBS twice taking care not to disrupt/detach cells and fresh PBS (100 µL) was placed in the wells and fluorescence was measured using a TECAN plate reader at 485 nm excitation and 590 nm emission. Cells were then placed in the incubator with fresh growth media in the wells and wash steps and fluorescence read at 15 min intervals for 2 h. Data for treated wells were normalised to vehicle controls. For pump inhibitor treatment, drugs were administered to the cells 30 min before epirubicin loading (verapamil - 20 µM, MK571 – 50 µM, and KO143 – 15 µM). All experiments were conducted in technical triplicate and are presented as mean ± SEM of three or more independent replicates.

### In vivo model

All protocols involving mice were approved by the Illinois Institutional Animal Care and Use Committee at the University of Illinois. Mouse 4T1 BCa cells were maintained in DMEM supplemented with 10 % calf serum. 1×10^6^ cells in a 1:1 ratio of PBS:matrigel were grafted orthotopic into the axial mammary fat pad of BALB/c mice. Mice were split into 4 groups (ten mice per group) of placebo, GW3965, epirubicin, or GW3965 and epirubicin combined. Daily treatment with GW3965 (30mg/kg) or placebo started one day post-graft. Treatment with epirubicin started two days post-graft and was administered every other day at 2.5mg/kg. Subsequent tumor growth was determined by direct caliper measurement. At the end of the experiment, mice were humanely euthanized, and tumors were dissected and weighed. Quantification of mRNA was carried out as described previously (Shahoei *et al*, 2019). Relative expression was determined via the 2-ΔΔCT method and normalized to housekeeping gene (TBP).

### Human samples: Ethical approval, collection, and processing

37 fresh/frozen tumour samples for LC-MS/MS and gene expression analysis were obtained with ethical approval from the from the Leeds Breast Research Tissue Bank (15/HY/0025). Selection criteria were all available ER-negative tumours with >3yr follow-up with fresh-frozen tumour material available. Supplementary Table 1 summarizes the clinic-pathological features of the ‘fresh/frozen’ cohort. For the tissue micro-array (TMA) 148 tumour samples were obtained with ethical approval from Leeds (East) REC (06/Q1206/180, 09/H1306/108). The patient cohort consisted of triple negative/ basal tumours as determined through immunohistochemistry.

Further selection criteria were that selected tumours had not undergone neoadjuvant therapy, contained sufficient tumour stroma, had low levels of inflammatory cells or necrotic material, and whether normal tissue was available from archival resection blocks (n=148). Suitable tumour areas were identified through haematoxylin/eosin stain and 0.6mm cores of tumour tissue removed in triplicate and deposited into recipient wax block. Supplementary Table 2 summarizes the clinic-pathological features of the TMA cohort.

### Statistics

Analysis of colony forming assays was performed using paired t-tests for comparisons between epirubicin treated cells with and without pre-treatment with LXR ligands, or one-way ANOVA after correction for multiple testing when comparing all treatments at once. Analysis of gene and protein correlations were assessed using Spearman’s correlation with linear regression.

Significance in gene expression analyses of genes implicated in chemotherapy resistance pre and post gene silencing and inhibitor loading was assessed using 2-way ANOVA. The half-life of intra-cellular epirubicin signal was determined using dissociation one phase exponential decay and the expression of Pgp in patient tumours that had suffered an event compared with those who had not was assessed using Mann-Whitney U tests. Patient survival was assessed using log rank tests and *in vivo* mouse experiments were assessed using 1-way ANOVA and corrected for multiple comparisons.

## Results

### LXR ligands influence epirubicin response in TNBC cells in vitro

To establish if the scOHC-LXR axis influences TNBC chemotherapy response, *in vitro* assays were used to test how combinations of epirubicin and LXR ligands altered tumor cell survival and growth. MTT assays were used as a readout of cellular health. LXR agonist treatment generally impaired epirubicin cytotoxicity, while antagonism of LXR enhanced efficacy. Apply the LXR synthetic agonist GW3965 increased the concentration of epirubicin needed to induce cell death (p<0.0001 for 0.1μM, 0.25μM and 1μM in MDA.MB.468 and p<0.01 for all concentrations in MDA.MB.231), as did the endogenous agonists 24OHC (p<0.05) for all concentrations in both cell lines, and 10μM 26OHC (also known as 27-hydroxycholesterol, the product of CYP27A1; p<0.05 for both cell lines), (Fig 1A). For all concentrations of the LXR antagonist GSK2033 tested (Fig 1B) in MDA.MB.468 cells there was a significant increase in epirubicin effect (p<0.0001). In MDA.MB.231 cells only 250nM was effective (p<0.05). Colony forming assays (CFA) were then conducted with cells pre-treated with LXR ligands or vehicle control, before being exposed to epirubicin. As expected, epirubicin treatment alone (25 nM for 24 h) resulted in a significant attenuation (p<0.001) of between 40-70 % of the cells’ ability to form colonies in the following days (note y axis for epirubicin in Fig 1C). When LXR agonists GW3965, 24OHC, or 26OHC were combined with epirubicin treatment (Fig 1C) we observed significant rescue of colony formation compared to the epirubicin alone (p<0.01 for all agonists in both MDA.MB.231 and MDA.MB.468). When the synthetic LXR antagonist GSK2033 was used however (Fig 1D) the reverse was observed and the efficacy of epirubicin was enhanced (p<0.05). All concentrations of LXR ligands tested could transactivate LXR (Hutchinson *et al*., 2019). Low doses that had no effect on proliferation (SF1A) or colony formation (SF1B), as well as higher doses that were antiproliferative in MTT were evaluated to avoid confounding chemoresistance with anti-proliferative effects. From these data we concluded that synthetic and endogenous LXR ligands conferred resistance to epirubicin.

**Figure 1:**
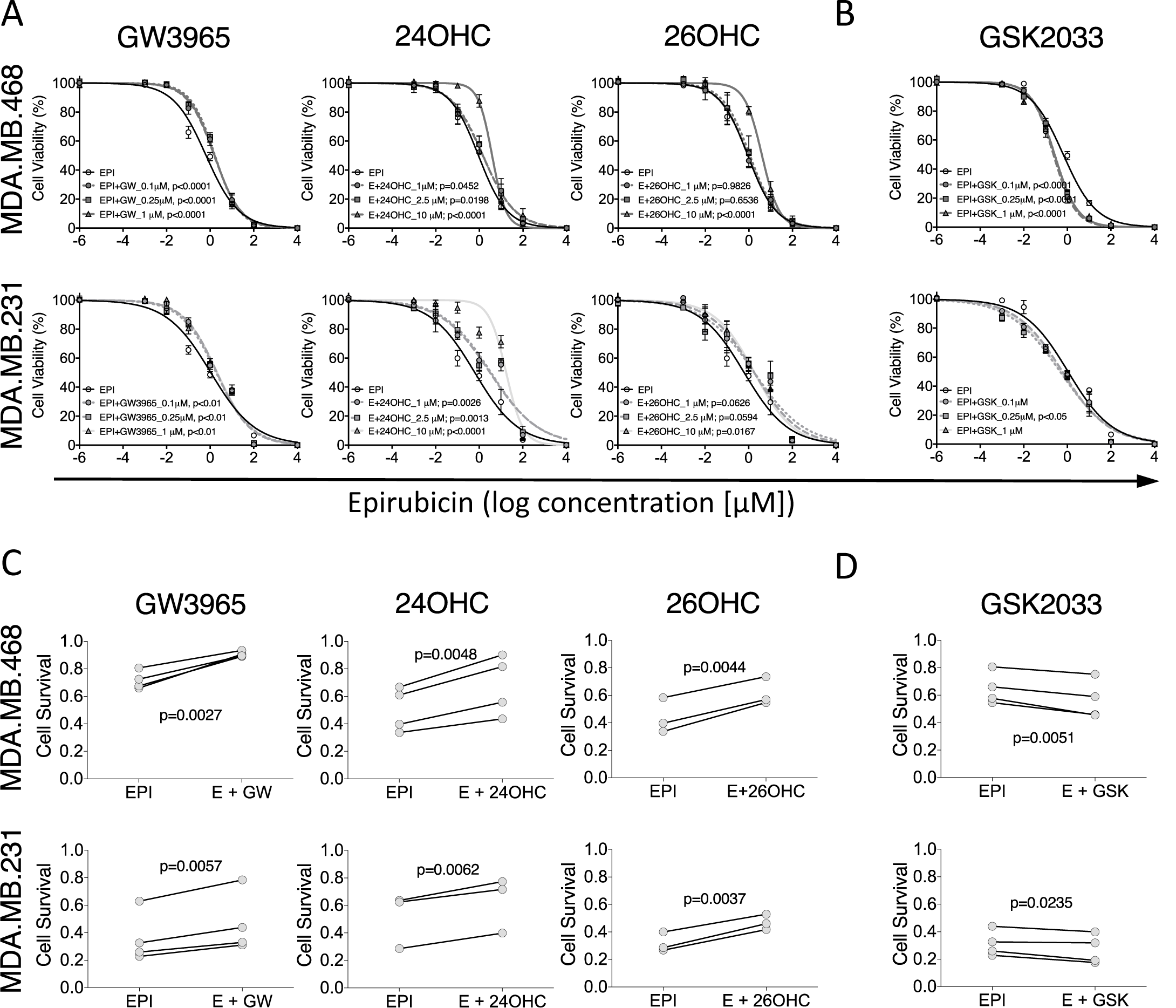
LXR ligands induce chemoresistance in triple negative breast cancer cell lines. MTT assay of epirubicin dose response following pre-treatment of MDA.MB.468 and MDA.MB.231 cell lines with low, medium, and high doses of LXR agonists (A) or antagonist (B). Data shown are mean with SEM of 3 independent replicates each performed with 6 technical replicates. Statistical significance was determined using non-linear regression curve comparison against epirubicin (EPI) only curve. Colony Forming Assays measuring how the ability of MDA.MB.468 and MDA.MB.231 cell lines to generate colonies after epirubicin exposure is improved by pre-treating cells with LXR agonists (C) or antagonist (D). Three or 4 independent replicates were performed (as indicated), each independent replicate is the mean of 3 technical repeats. p-values were calculated using paired t-test.

### LXRα is linked to Pgp expression and function in TNBC patients and in vitro

The chemotherapy efflux pump Pgp was a strong candidate as the mediator of this LXR ligand induced resistance to epirubicin, as it has previously been reported to be modulated by oxysterols in the blood brain barrier (Saint-Pol *et al*., 2013). To establish if Pgp is potentially regulated by LXRα or LXRβ, and thus sensitive to cholesterol metabolic flux in the tumor, we mined publicly available transcriptomics datasets from TNBC tumours to determine correlation of mRNA expression, and binding of the LXRs to the Pgp promoter.

Expression of LXRα mRNA was weakly, although significantly, correlated with Pgp (Fig 2A) in both METABRIC (n=313; p=0.0024; R=0.17) and TCGA datasets (n=95; p=0.0076; R=0.27). No correlations between LXRβ and Pgp were observed (p>0.05). Other drug efflux pumps (MRP1 and BCRP) were not correlated with LXRα in either dataset (SF2; p>0.05). We next investigated if LXRα or LXRβ could directly bind the Pgp promoter using mouse macrophage and liver tissue datasets, available in the cistrome.org ChIP-Seq repository. LXRα, but not LXRβ, accumulated in the promoter region of Pgp in response to LXR agonist (SF3). As a positive control, we also confirmed that both LXRα and LXRβ accumulated in the promoters of known LXR target genes ABCA1 and APOE (SF3). These data are consistent with the hypothesis that LXRα, but not LXRβ, is able to bind the Pgp promoter region. Furthermore, expression of LXRα, but not LXRβ, is positively correlated with Pgp expression in the tumours of TNBC patients in separate publicly available datasets.

**Figure 2:**
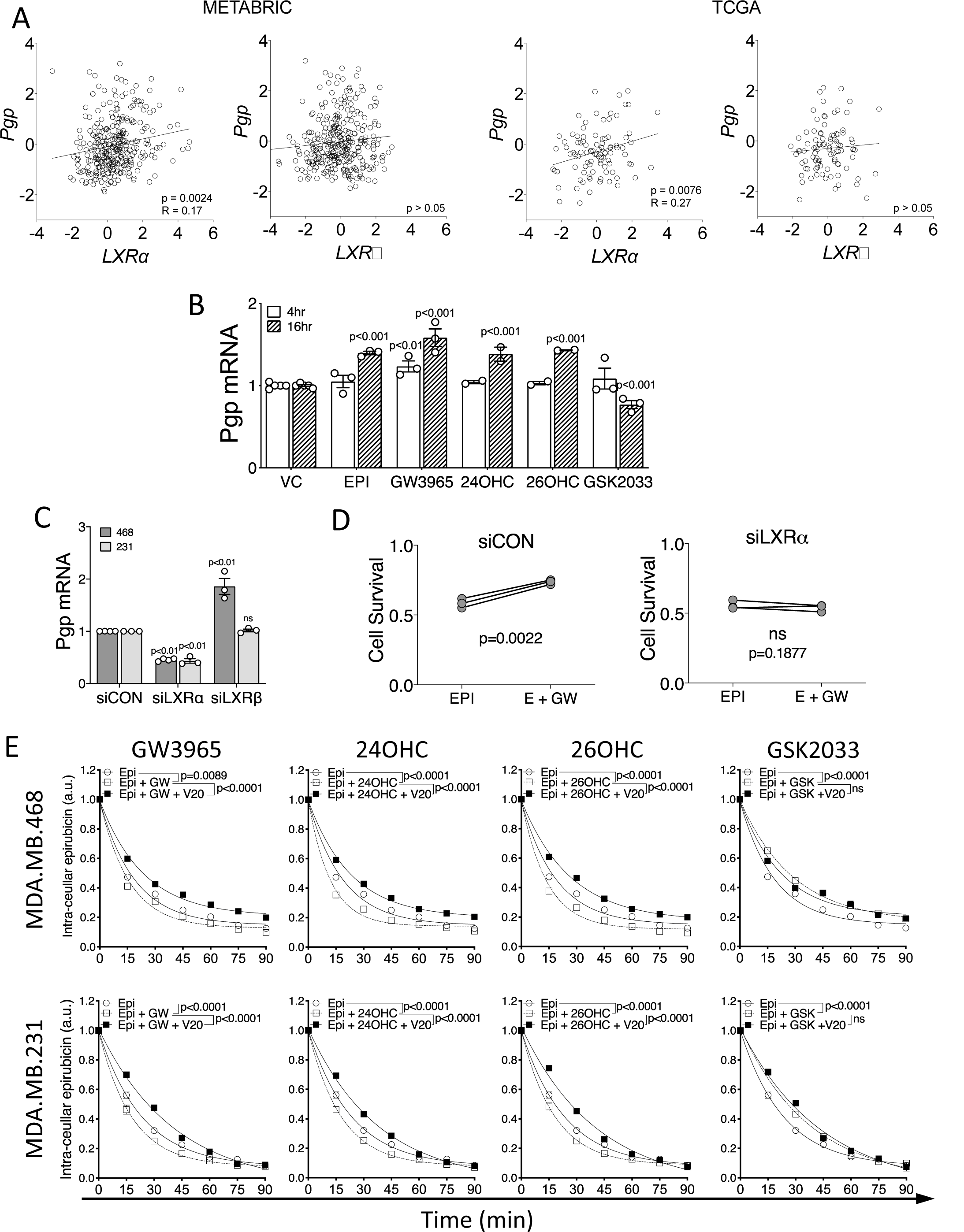
Pgp expression and function is linked to LXR activity. Correlation analysis in TNBC tumors of *LXR*α or *LXR*β with *ABCA1* and *Pgp* gene expression obtained from obtained from METABRIC (n=313) and TCGA (n=95) datasets accessed via cBioportal (A). Statistical significance was assessed using Pearson’s correlation test with linear regression. Pgp expression measured at indicated time points after ligand exposure (B) or siRNA knockdown of LXRα or LXRβ (C). B-C Symbols show independent replicates and bars show mean with SEM. Statistical analysis was performed using 2-way ANOVA and is representative of 3 independent replicates with SEM. Colony forming assay in MDA.MB.468 cells after loss of LXRα by siRNA showing GW3965 (GW) no longer protects cells from epirubicin in the absence of LXRα (D). MDA.MB.468 and MDA.MB.231 cells were pre-treated with LXR ligands or vehicle for 16 h and Pgp inhibitor (verapamil 20 μM [V20]) or vehicle for 30 min, before loading with epirubicin (50 μM) for 1h. The half-life of the intra-cellular epirubicin signal was calculated using dissociation one phase exponential decay. NB: epirubicin curve data are replicated in each graph for visualization purposes. Data shown are mean with SEM of 3 or 4 independent replicates each made from six technical replicates. Significance was calculated using non-linear regression curve comparison against epirubicin (EPI) only curve.

To ascertain if Pgp was regulatable by LXR ligands in TNBC cells, we treated MDA.MB.468 and MDA.MB.231 cells with a range of LXR ligands and found significant up- and down-regulation of Pgp in response to agonists or antagonists respectively at two different time-points (Fig 2B). To test which LXR isoform was primarily responsible for Pgp induction, siRNA was used to knock down LXRα and LXRβ separately. siLXRα significantly diminished Pgp expression (Fig 2C; p<0.01), demonstrating the LXRα positively regulates Pgp, and also halted the agonist GW3965’s ability to protect cells from epirubicin in CFA (Fig 2D; p>0.05), demonstrating the functional impact of LXRα’s control of Pgp. Knock-down of LXRβ did not reduce Pgp expression. siLXRα and siLXRβ knockdown efficiency was confirmed at protein (SF4A) and mRNA levels (SF4B), as was loss of expression of canonical target gene expression (SF4B). Knock down of either LXR isoform had no effect on expression of MRP1 (SF4C; p<0.05) or BCRP efflux pumps (SF4D; p<0.05).

After establishing Pgp expression was at least in part dependent on the scOHC-LXRα axis, and that this signaling pathway protected TNBC cells from chemotherapy, we hypothesized that Pgp’s drug efflux function was crucial. We developed a novel chemotherapy efflux assay allowing time-resolved monitoring of intracellular epirubicin concentration in a high throughput system by exploiting epirubicin’s natural fluorescence. After pre-treating MDA.MB.468 or MDA.MB.231 cells with vehicle control (VC; ethanol), LXR agonist (GW3965, 24OHC, or 26OHC) or antagonist (GSK2033) cells were then ‘loaded’ with epirubicin and intra-cellular epirubicin measured at 15 min intervals as the epirubicin was pumped out. The half-life (*t*_1/2_) of epirubicin signal in vehicle treated control cells was 13.05 min for MDA.MB.468 cells and 16 min for MDA.MB.231 (Fig 2E) and all pre-treatments were compared back to this vehicle/epirubicin condition with one-phase decay non-linear regression model. GW3965 pre-treatment led to a significant reduction in epirubicin retention time to *t*_1/2_ = 10.98 min (p=0.00899) and *t*_1/2_ = 12.28 min (p<0.0001) in MDA.MB.468 and MDA.MB.231 cells respectively (note shift of Epi curve [open circles with solid line] to left with addition of GW3065 [open squares with dotted line]). scOHC pre-treatment also significantly (p<0.0001 for all) shifted curves to the left, again indicating decreased epirubicin retention time (468 cells: 24OHC *t*_1/2_ = 8.27 min, 26OHC *t*_1/2_ = 9.62 min; 231 cells: 24OHC *t*_1/2_ = 12.28 min, 26OHC *t*_1/2_ = 12.38 min). This enhanced efflux of epirubicin was reversable by the Pgp specific inhibitor verapamil (V20) (note shift of curve to the right with V20 as denoted by closed squares), demonstrating the dependence on Pgp function. In contrast, inhibiting LXR activity with the synthetic antagonist GSK2033 led to significantly increased retention of epirubicin (468 cells: *t*_1/2_ = 20.19 min; 231 cells: 231 cells *t*_1/2_ = 25.86 min p=0.0004), and this was not reversed or enhanced by verapamil (p>0.05). Overall intra-cellular epirubicin accumulation was also lower in LXR agonist treated cells (SF5A). Experiments with selective inhibitors demonstrated that Pgp but not other drug efflux pumps (MRP1, BCRP) was necessary and sufficient for LXR mediated epirubicin efflux in control experiments (SF5B-C).

### LXR activation confers chemoresistance and enhances Pgp expression in vivo

To corroborate *in vivo* these chemoprotective effects of the scOHC-LXR axis, a preclinical model using TNBC syngeneic 4T1 cells grafted in mammary fat pads of Balb/c mice was evaluated. Mice in four groups were treated with either vehicle control, GW3965, epirubicin, or GW3965 and epirubicin combined. As expected, epirubicin reduced tumour growth (Fig 3A) and final size (Fig 3B), as did GW3965. The tumors of mice in the combination group grew significantly more quickly (p<0.0001) and were significantly larger (p<0.0001) than the epirubicin alone group. Analysis of resected tumors indicated significantly increased expression of both canonical *Abca1* (p<0.001) and *Abcb1b* (the mouse gene for Pgp) (p<0.0001) in GW3965 treated animals compared to controls (Fig 3B), demonstrating that the LXR agonist had effectively activated relevant target genes within the tumour tissue. Mice in all groups gained weight at a similar rate through the experiment (p>0.05) indicating mouse health was similar between groups over the experimental period (SF6A). Other drug efflux pumps were not found to induced by GW3965 in this model (SF6B). We concluded that activation of LXR *in vivo* induced expression of Pgp and confers chemoresistance as observed *in vitro*.

**Figure 3:**
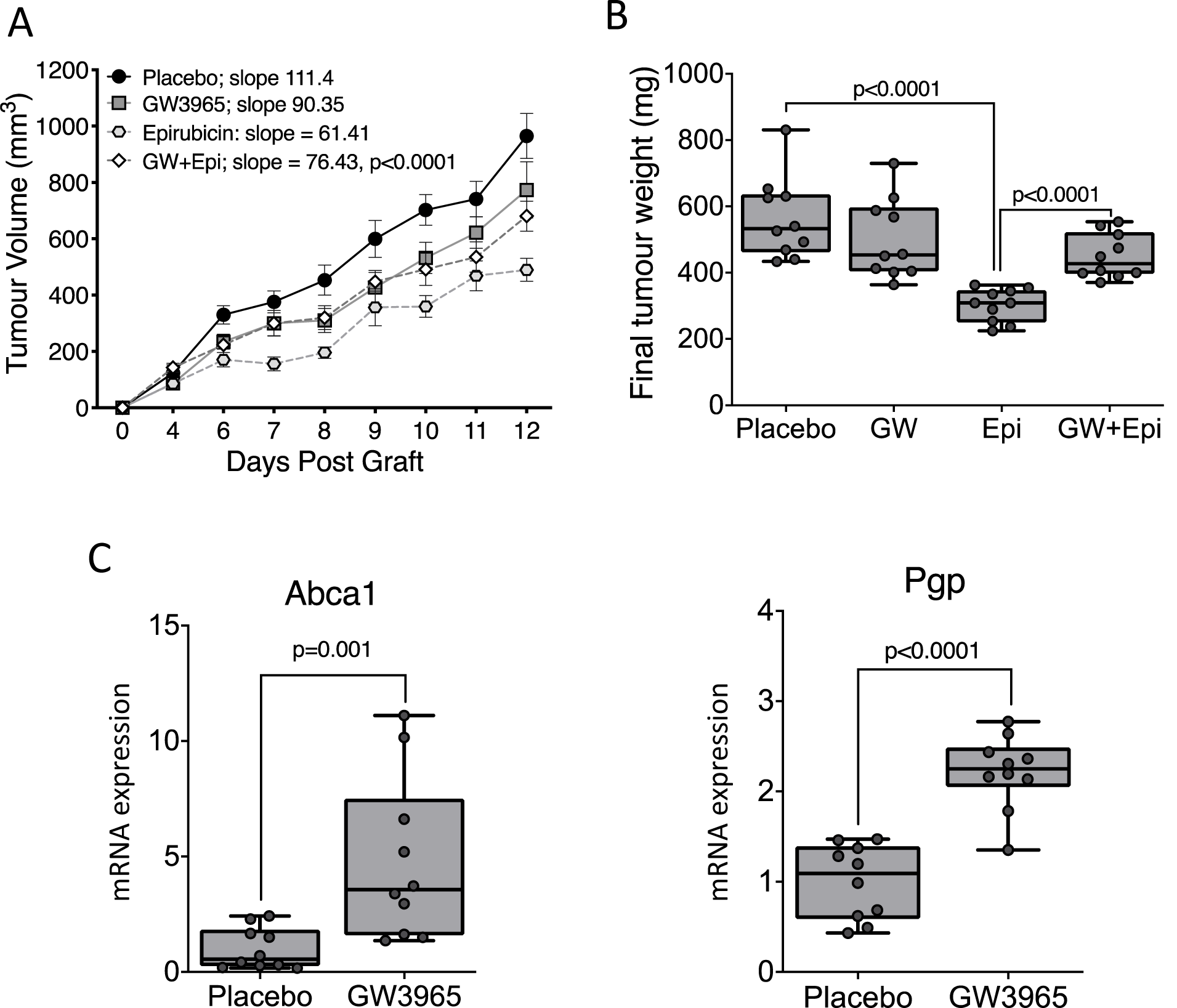
LXR activation drives Pgp expression and chemotherapy resistance in vivo. 4T1 cells (TNBC) were grafted orthotopically into the axial mammary fat pad of BALB/C mice. Mice were treated with either placebo or the LXR ligand GW3965 (daily, 30 mg/kg) 24 h post-graft. Treatments with placebo or epirubicin (every other day, 2.5 mg/kg) commenced 48 h post-graft. Tumor volumes measured by calipers (A) or tumor weight (mg) after 12 days (B). Statistical analysis was assessed using non-linear regression. Expression of Abca1, and Pgp was assessed by qPCR analysis (C). Statistical analysis was assessed using one way ANOVA with SNK test, with 10 mice per group (individual symbols) and shown with median and range.

### LXR and its ligand regulators are correlated with Pgp and are prognostic indicators in ER-negative BCa patients

To investigate if LXRα protein was associated with Pgp expression in ER-negative breast cancer patients, ABCA1 mRNA, Pgp mRNA, LXRα protein expression were measured in a cohort of ER-negative tumor samples from the Leeds Breast Research Tissue Bank (LBRTB cohort: n=47; patient characteristics reported in Supplementary Table 1; representative immunoblots shown in SF7).

LXRα protein was positively correlated with mRNA expression of Pgp (Fig 4A; p=0.0046; r=0.43), validating previous mRNA analyses from public datasets (Fig 2A). Kaplan-Meier survival analyses after the cohort was dichotomized into low and high expression groups for LXRα indicated that high LXRα (Fig 4B; p=0.0051) were associated with significantly worse prognosis.

**Figure 4:**
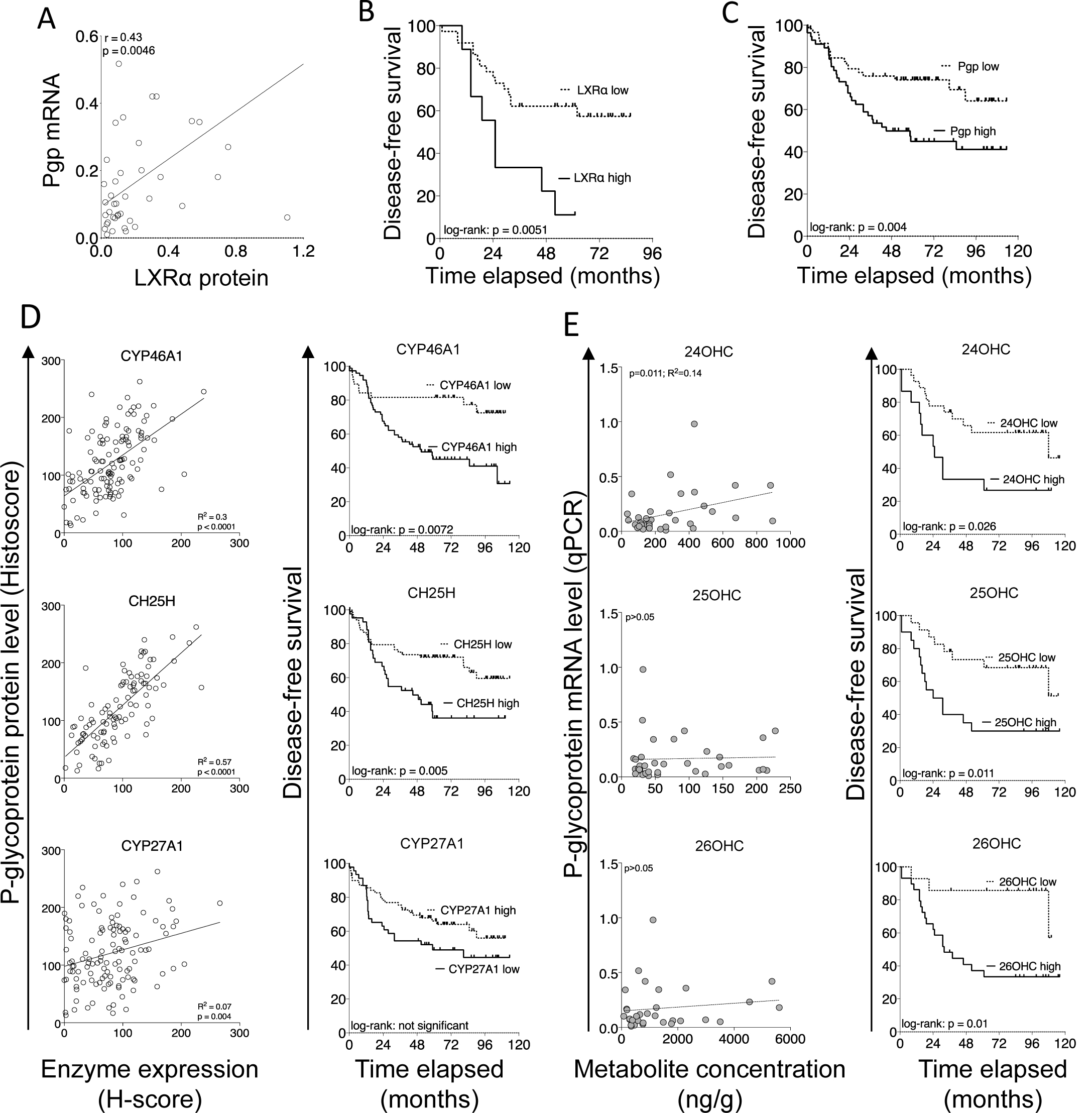
LXRα protein expression is correlated with Pgp expression and is a molecular marker of poor prognosis in ER-negative breast cancer patients. Correlation analysis between LXR protein expression and matched mRNA of Pgp in 47 ER-negative breast tumors (relative to HPRT). Line indicates linear regression, correlations calculated using Pearson correlation. Kaplan Meier survival plots following stratification of the cohort into high and low LXRα (B) and Pgp (C) expression. Log-rank tests used to calculate significance. (D) Tissue microarray of 148 TNBC tumors stained for indicated proteins determined by immunohistochemistry and correlated with Pgp protein (left) and disease-free survival (right). (E) scOHC concentrations determined by LC-MS/MS and correlated with Pgp protein (left) and disease-free survival (right). Correlations calculated with Pearson’s test and survival differences with log-rank test and Kaplan-Meier curves.

To explore if Pgp protein expression and patient prognosis were associated with synthesis of OHC, we undertook two analyses. First, a large-scale analysis of scOHC synthesizing enzyme expression in TNBC tumors in a microarray of formalin fixed tissue (TMA cohort: n=148; patient characteristics reported in Supplementary Table 2). Pgp protein expression in TNBC epithelial cells was a strong indicator or shorter disease free survival (Fig 4C; p=0.004), and expression was strongly and significantly positively correlated with protein expression CYP46A1 (p<0.0001; R^2^=0.3) and CH25H (p<0.0001; R^2^ = 0.57), and weakly with CYP27A1 (p=0.004; R^2^=0.07) (Fig 4D – left panels), enzymes that synthesize 24OHC, 25OHC, and 26OHC respectively.

Shorter disease-free survival was found in patients with high expression of CYP46A1 (p=0.005) and CH25H (p=0.0072), but not CYP27A1 (p>0.05) (Fig 4D - right panels). Secondly, a smaller analysis was performed in the LBRTB cohort where fresh-frozen tissue was available thus allowing direct LC-MS/MS measurements of scOHC concentrations in matched tumour samples to protein lysates measured in Fig 4D. 24OHC (but not 25OHC or 26OHC) metabolite concentrations were significantly positively correlated with Pgp mRNA expression (Fig 4E – left panels); high concentration of all three metabolites (24OHC p=0.026; 25OHC p=0.011; 26OHC p=0.01) were prognostic of shorter disease-free survival (Fig 4E – right panels). From these data we concluded that multiple components of the LXR signaling pathway were likely to converge on Pgp mediated drug resistance and contribute to shorter disease-free survival in TNBC patients.

### LXRα and its ligands are decoupled from Pgp in surviving TNBC patients, but remain linked in patients who suffer relapse or die

After establishing a mechanism linking the cholesterol metabolic sensor, LXRα, with chemotherapy resistance at the molecular level, and discovering a discernable effect on survival in cell lines, pre-clinical models and patients, we wanted to explore if measuring activity of the scOHC:LXR pathway could separate surviving patients from those who relapse or died from their disease. We split the LBRTB cohort (Supplementary Table 1) into a ‘no event’ group, i.e. those who did not have a recurrence during follow up (median follow up time = 96 months), and an ‘event’ group, i.e. those who had either died from their disease or had relapsed (median time to event = 20 months). Protein expression of LXRα (p=0.076), and mRNA expression of ABCA1 (p=0.0121) and Pgp (p=0.0015) were higher in the ‘event’ group (Fig 5A) indicating greater activity of the LXR signaling axis in the event group; interestingly, the concentrations of scOHC metabolites were not significantly different between the event and no event groups (SF8). We also tested the correlations between expression of Pgp and its regulators LXRα and 24OHC in the cohort, separating patients who had an event from those who didn’t; LXRα protein was only correlated with Pgp in the event group (Fig 5B; p=0.0012) and similarly, 24OHC was only correlated with Pgp in the event group (Fig 5C; p=0.01). We tested this relationship in the METABRIC cohort and found that LXRα only correlated with Pgp in patients who had died from their disease or who had relapsed (SF9; p=0.017).

**Figure 5:**
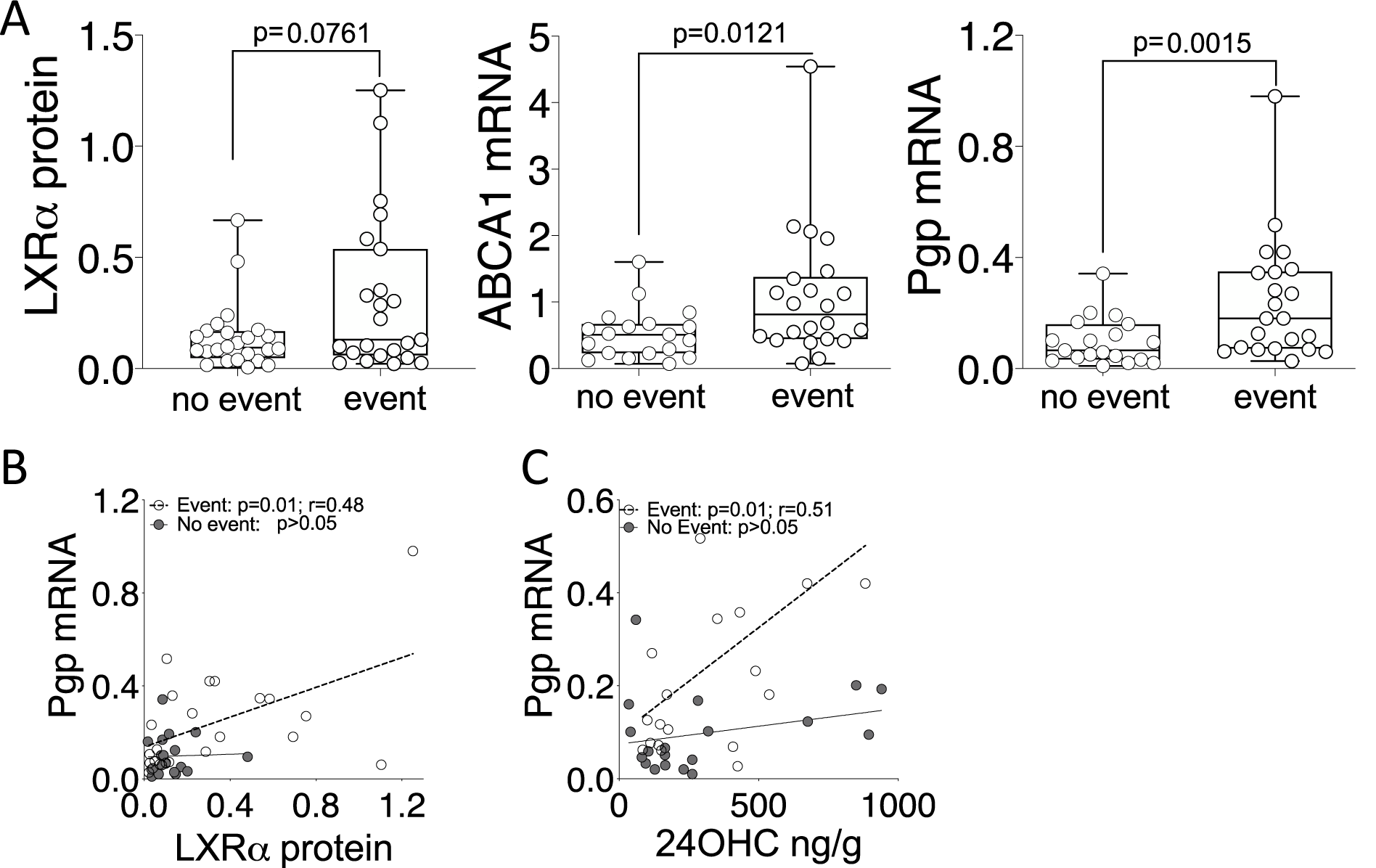
LXR is decoupled from Pgp in the tumours of patients who survive. Protein expression (LXRα), and mRNA expression for ABCA1 and Pgp (A) were assessed using one-tailed Mann-Whitney test in the LBRTB cohort comparing event and no event groups. Pearson correlation analysis between matched Pgp mRNA and either immunoblotting for LXRα protein (B), LC-MS/MS for 24OHC (C).

## Discussion

The purpose of this study was to evaluate the role of cholesterol side-chain hydroxylation products on chemotherapy resistance, specifically in TNBCs – the breast cancer subtype in which chemoresistance is most problematic. We demonstrated that LXRα activation is linked to chemotherapy resistance *in vitro* and *in vivo*, and to worse patient survival. We conclude that this is due, at least in part, to the chemotherapy efflux pump Pgp being a transcriptional target of the scOHC:LXR axis, thus linking cholesterol status with the innate ability of a triple negative tumor cells to evade chemotherapy. A major observation from our data was that patients stratified by survival status had divergent LXR signalling pathways; the tumours of the patients who died were enriched for scOHC synthesizing enzymes and the metabolite 24OHC, and there was a positive correlation between Pgp and individual components of the signaling axis. In patients who did not have recurrences, these signaling components were not correlated with Pgp, irrespective of which cohort we evaluated, indicating that actual decoupling of LXR from Pgp in these tumours was contributing to survival.

The role of drug efflux pumps such as Pgp, BCRP, and MRP1 in TNBC resistance have been described (Kim *et al*, 2013; Kim *et al*, 2015). Yet directly targeting these pumps has largely failed as a clinical strategy due to the essential roles that xenobiotic efflux pumps play in healthy tissue, from side effects from drug-drug interactions, and lack of specificity (Yang *et al*, 2018). The factors that determine whether pCR is achieved are important to elucidate because patients for whom only a partial pathological response with NACT is achieved typically do worse than those who elect for ACT (Xia *et al*., 2020). Cancer specific mechanisms of efflux pump regulation have been identified, but until now have largely been attributed to oncogenic signalling pathways converging on their transcriptional regulation (Das *et al*, 2013; Kim *et al*., 2015). The data reported here however indicate a new direction; we provide evidence of a direct link between the expression and function of Pgp, and metabolic flux of cholesterol, a nutrient that is modifiable by diet, lifestyle, and existing pharmacological approaches. In this context, the findings presented here lead to speculation that two clinical hypotheses are appropriate for further evaluation. Firstly, patients stratified based on LXR activity may be predicted to achieve different rates of pCR after NACT, or survival after ACT. Secondly, a therapeutic adjunct that limits scOHC synthesis may improve pCR rates. Cholesterol lowering approaches are already in clinical practice for the management of cholesterol related diseases. Excitingly, these interventions could be tailored to the patient to offer an element of control to individuals who may otherwise seek lifestyle an dietary guidance from sources that have not been peer reviewed (Thorne *et al*, 2020). It of course remains to be determined if such strategies would be beneficial in improving survival rates via chemosensitization, but it is encouraging that extra-hepatic scOHC levels are modifiable by diet (Guillemot-Legris *et al*, 2016; Sozen *et al*, 2018) as well as circulating levels being highly responsive to statins (Dias *et al*, 2018). A meta-analysis of statin use in unstratified breast cancer patients has shown inhibition of cholesterol synthesis was associated with protection from relapse and death in the first 4 years post-diagnosis (Liu *et al*, 2017a), which interestingly, is the period of highest relapse risk for TNBC patients (Liedtke *et al*., 2008).

LXR ligands are being explored in clinical trials as novel therapeutics, and as diagnostic and prognostic tools. Within these studies, important distinctions between activation of LXRα and LXRβ need consideration. The histamine conjugated oxysterol such as dendrogenin A, preferentially activates LXRβ and drives lethal autophagy (Poirot & Silvente-Poirot, 2018) and differentiation of breast cancer cells (Bauriaud-Mallet *et al*., 2019), and synthetic the LXRβ agonist RGX-104 enhances cytotoxic T lymphocyte tumour destruction (Tavazoie *et al*, 2018). Dual LXRα/LXRβ ligands such as 26OHC convincingly slow tumour growth through the anti-proliferative activities of LXR, drive LXR dependent metastasis (Nelson *et al*, 2013), and substitute for estrogen to drive ER dependent breast cancer growth (Nelson *et al*., 2013; Wu *et al*, 2013). Furthermore, circulating dual LXR oxysterol ligands may be prognostic and therapeutic indicators (Dalenc *et al*., 2017). Our results suggest efficacy of cytotoxic therapies may be reduced if LXRα is stimulated; caution should be applied in clinical application of dual LXR ligands or of LXRα agonists, as novel cancer therapeutics.

Mechanisms that lead to innate chemoresistance have previously been flagged as a critical research gap (Eccles *et al*., 2013). LXR is a sensor and homeostatic regulator of nutritional status, so our data support the hypothesis that a cholesterol rich environment during tumor development may actually contribute to innate chemoresistance, thus addressing a critical research gap. Ligand dependent activation of LXR in many tissues is controlled by factors that can be influenced by the diet and existing pharmacological agents, making this pathway attractive for therapeutic targeting. Encouragingly, further clinical investigation where dietary advice or statins that acutely lower circulating or tissue scOHC levels could evaluate LXR modulation to improve pCR rates in this hard to cure subgroup of breast cancer patients. In conclusion, endogenous synthesis of LXR ligands and LXRα activity are prognostic indicators for TNBC patient survival and should be explored prospectively in the clinical setting as functional and targetable prognostic indicators for response to chemotherapy.

