## Supplementary Information for "Liver x receptor alpha drives chemoresistance in response to side-chain hydroxycholesterols in triple negative breast cancer"

**Supplementary Data**

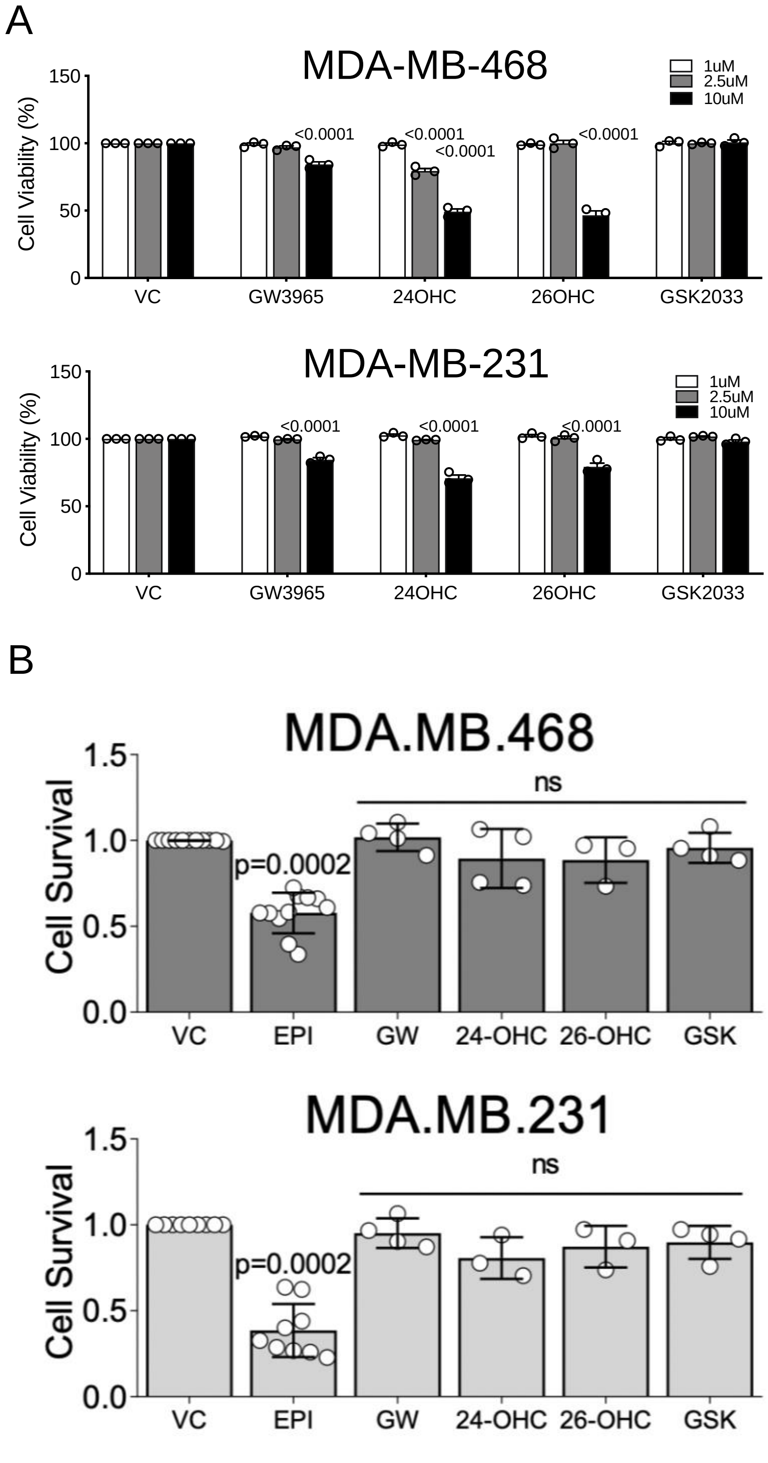


**Supplementary Figure S1: Effect of single treatments in MTT and CFA**

Effect of single ligand treatments on cell viability in MTT (A) and CFA (B). For OHC, concentrations were 1 μM (low), 2.5 μM (Medium), and 10μM (High). For synthetic ligands GW3965 and GSK2033 concentrations were 100nM (Low), 250nM (Medium), and 1μM (High). Viability reported here indicated the effect of pre-treatments only. Experiments were repeated in 3-5 independent replicates (indicated by circles). Statistical differences were determined using two-way ANOVA (A) or one-way ANOVA (B). ns = not significant. VC – vehicle control.

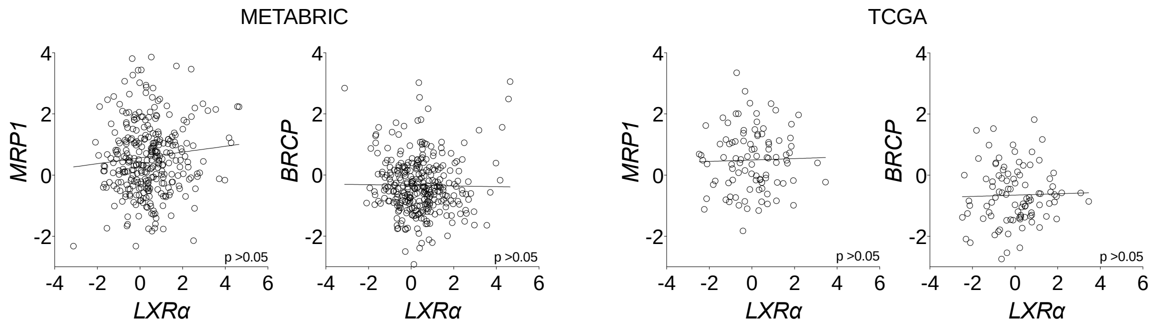


**Supplementary Figure S2: MRP1 and BCRP do not correlate with *LXRα* in TNBC**

Correlation analysis in TNBC tumors of *LXR*α with chemotherapy resistance pumps *MRP1/ABCC1* and *BCRP/ABCG2* gene expression obtained from obtained from METABRIC (n=313) and TCGA (n=95) datasets accessed via cbioportal. Statistical significance was assessed using Pearson's correlation.

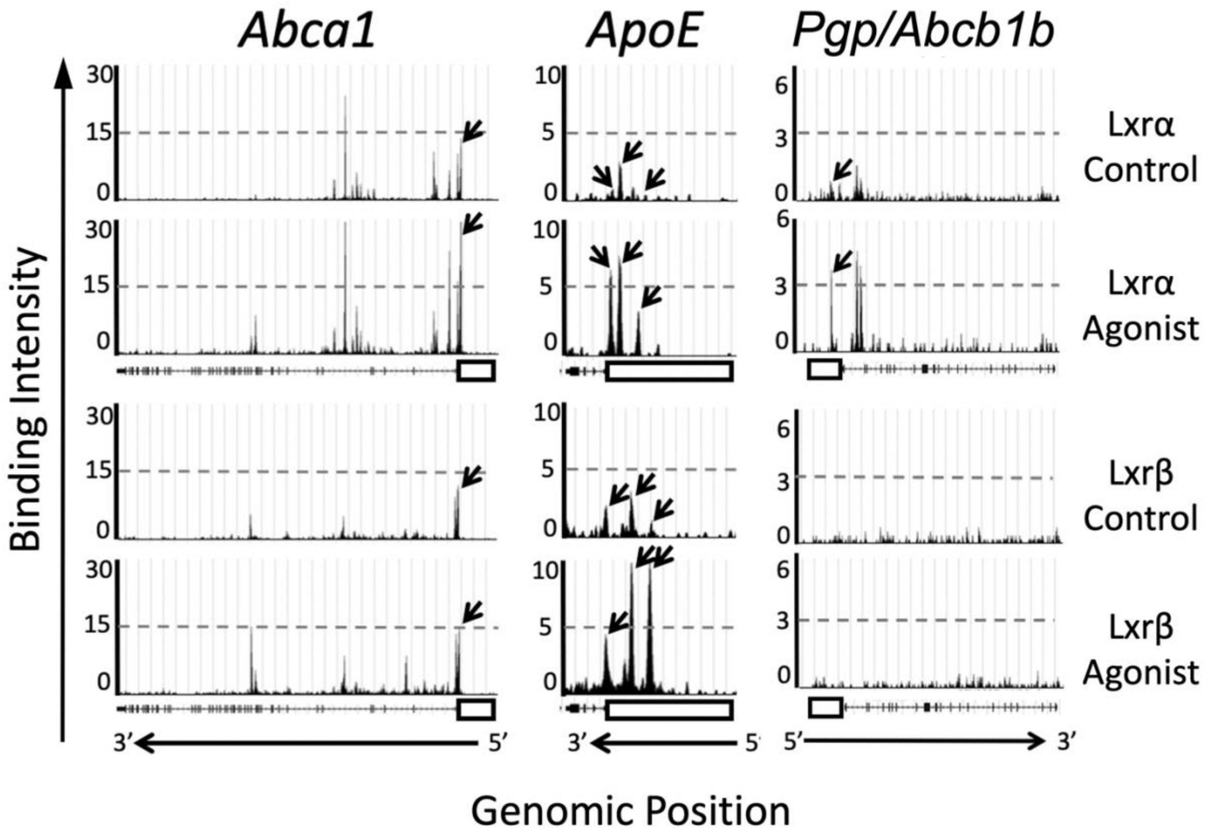


**Supplementary Figure S3: Binding of LXR to target gene promoters**

Lxrα binding was assessed in a ChIP-Seq mouse macrophage dataset by (Oishi *et al*, 2017) with no treatment (ID: 72545) and after GW3965 treatment for 24 h (ID: 72544). Lxrβ binding was assessed in a ChIP-Seq hepatocyte datasets (Boergesen *et al*, 2012) with no treatment (ID: 5415) and after T0901317 treatment daily for 14 days (ID: 5416). 10kB of the promoter regions are displayed (open box) for *Abca1*, *Apoe* and *Pgp/Abcb1b.*

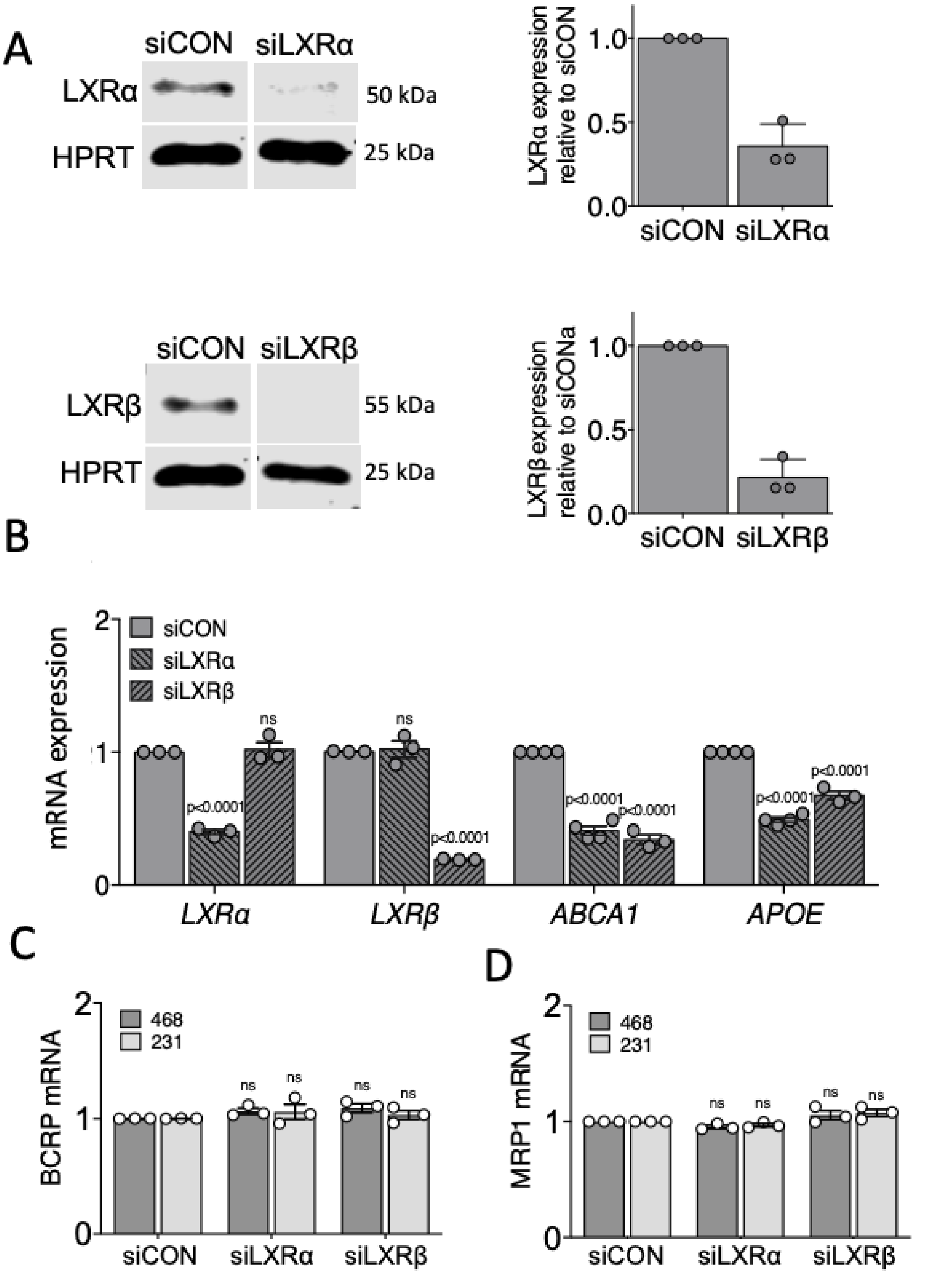


**Supplementary Figure S4: Validation of siLXR specificity and sensitivity**

The expression of the two LXR isoforms LXRα and LXRβ was knocked down using siRNA, and protein (A) and mRNA (A) were measured. Loss of expression of two LXR target genes ABCA1 and APOE was confirmed by qPCR. BCRP and MRP1 were not altered by siLXR knock-down. Statistical significance was established using 2-way ANOVA. Mean of 3-4 independent replicates (indicated as circles) with SEM are presented. Ns = not significant (p>0.05).

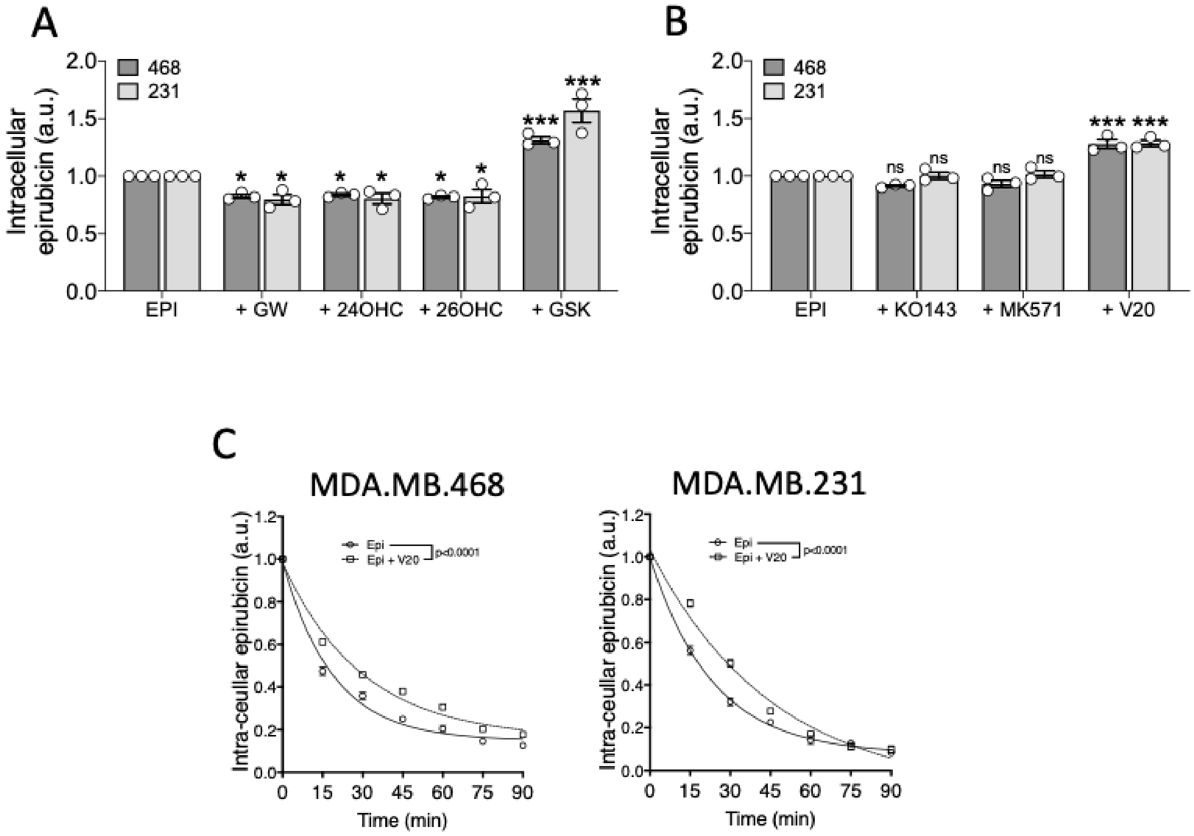


**Supplementary Figure S5: Cellular loading and efflux kinetics of epirubicin in TNBC cells**

MDA-MB-468 and MDA-MB-231 cells were seeded into black 96-well plates and after 24 h they were pre-treated with (A) indicated LXR ligand for 16 h or (B) ABC xenobiotic pump inhibitors KO143 (BCRP/ABCG2 inhibitor), MK571 (MRP1/ABCC1 inhibitor) or verapamil ([V20] p-gp inhibitor) for 30 min before a treatment of epirubicin (50 µM) for 1 h. Cells were washed with PBS and epirubicin loaded within the cells was measured fluorescently. Data shown are mean of three independent replicates (shown as circles) with SEM. 2-way ANOVA corrected for multiple testing was used for statistical testing. *≤ 0.05, **p<0.01, ***p<0.001, p<0.0001. (C) MDA-MB-468 and MDA-MB-231) cells were treated with a p-gp inhibitor (verapamil 20 µM [V20]) or vehicle for 30 min, before cells were loaded with epirubicin (50 µM) for 1h. The natural fluorescence of epirubicin was measured at 15 min intervals for 90 min. The half-life of the intra-cellular epirubicin signal was determined using dissociation one phase exponential decay, data shown are mean of 3-4 independent replicates with SEM.

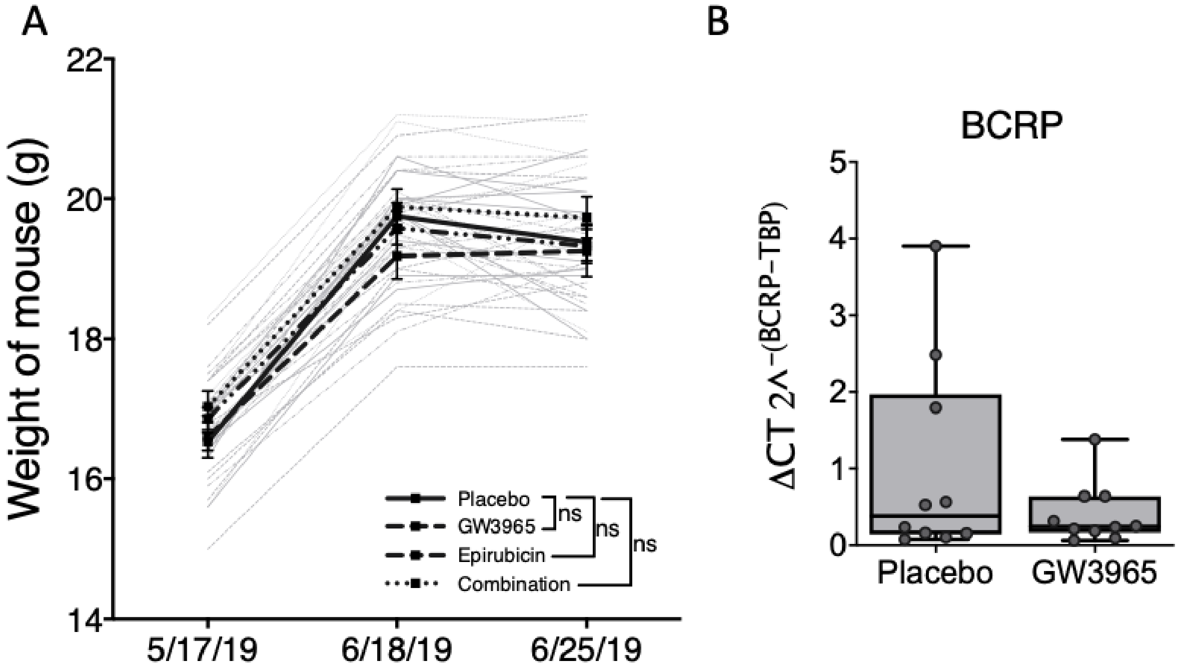


**Supplementary Figure S6: Mouse growth rate was not affected by treatment regimens**

40 mice were randomly allocated to four groups (10 per group). Mouse weight was recorded to ensure treatments did not affect growth rates. Individual mice are shown in gray, averages with SEM are shown in bold (A). *BCRP/ABCG2* mRNA was measured with qPCR in tumours excised from mice at the end of the experiment (B). Mann-Whitney U test was used to test for statistical differences, none were found.

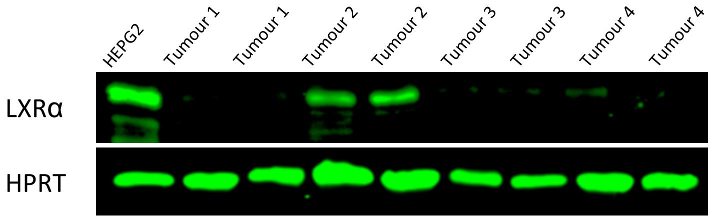


**Supplementary Figure S7: Representative immunoblots of LXRα and housekeeping control from 4 tumours**

Representative immunoblots for LXRα and LXRβ protein expression in four tumours (in duplicate lysate preparations) from the Leeds Breast Research Tissue Bank (A).

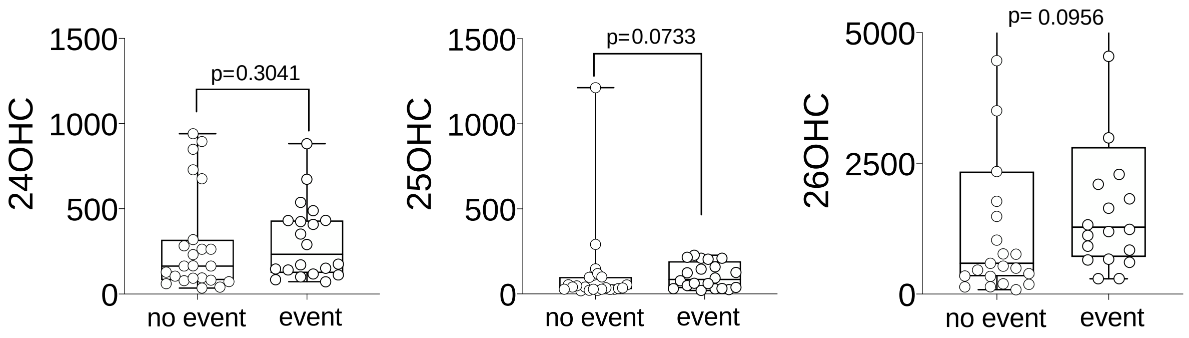


**Supplementary Figure S8: Intra-tumoral scOHC concentration is the same in patients who relapse compared to those who are disease free and alive**

Patients divided into those with who died from their disease or who relapsed (event: n=20) and those who are alive at the latest follow up (no event: n=23). A 2-tailed Mann-Whitney test was used to determined significant differences.

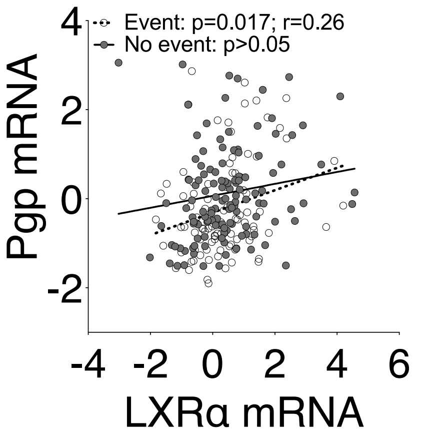


**Supplementary Figure 9: LXRα is positively correlated with Pgp in patients who relapse/die in METABRIC**

The correlation in event (n=92) and loss of correlation in no event (n=126) groups was confirmed in METABRIC at mRNA level.

### Supplementary Methods and Materials

#### Public dataset analyses

mRNA expression z-scores were obtained from METABRIC (Curtis *et al*, 2012) and TCGA (Cancer Genome Atlas, 2012) via cbioportal (Cerami *et al*, 2012). Analyses were performed as previously described (Hutchinson *et al*, 2019)Patients from both datasets were limited to *bona fide* ER-/PR-/Her2- using clinical annotation functions. Gene expression data were extracted and plotted in graphpad prism. To restrict by 'event' clinical status was set to died-from-disease or relapse, and 'no event' was alive or died-not-from-cancer. LXR ChIP-Seq by (Oishi *et al.*, 2017) and (Boergesen *et al.*, 2012) was identified in cistome.org (Liu *et al*, 2011) and selected following selection criteria for paired datasets (same organism, matched control and agonist treated) and quality control. Genes were visualized using the UCSC genome browser, aligned and cropped to include the gene of interest and 10 kb promoter region.

#### Antibody validation

For protein expression analysis, 150,000 MCF7 or HepG2 cells were seeded onto 22x22mm coverslips (631-0125, VWR International, UK), treated with siRNA as stated previously and fixed in 100uL of 3.7% paraformaldehyde for 15 minutes and washed twice in PBS at 72 following siRNA exposure Cells were incubated in PBS-Tween20 2% (PBS-T) for 30 minutes and blocked with 10% normal goat serum (NGS) (50197Z, Thermo Fisher Scientific, UK) for a further 30 minutes. Primary antibodies were added in 2% NGS in PBS-T and incubated for 1 hour at room temperature. Cells were washed twice in PBS-T and incubated for 30 minutes at room temperature with goat anti-rabbit antibody conjugated to alexa fluorophore 647 (Table 1) for 30 minutes. Coverslips were washed 4 times with PBS and mounted onto SuperFrost Plus slides (Menzel-Glaser; Braunschweig, Germany) with one drop of Prolong Gold Antifade Mountant with DAPI (P36941, Thermo Fisher Scientific, UK) and imaged using Zen-Imaging software (Zeiss, UK) on a LSM 880 with Airyscan (Zeiss, UK). Knockdown efficiencies were analysed through imaging three fields on each slide, drawing around the perimeter of all cells in the image and quantifying pixel intensity via the Mean Gray Value function in Fiji (LOCI, USA). siRNA validation of antibody specificity is shown (SF11).

#### Immunohistochemistry

IHC was performed as described previously with following modifications (Thorne *et al*, 2018). Antibodies validation using siRNA and immunofluorescence in cell lines with known high expression of each antigen is described in supplementary materials. Immunohistochemistry staining was quantified using histoscore as previously reported (Kim *et al*, 2013). Examples of negative, weak, moderate, and strong staining is given for each antibody (Fig SF12A). Tissue and TMA Blocks were sectioned to 5μM onto SuperFrost Plus slides (Menzel-Glaser; Braunschweig, Germany). Sections were dewaxed in xylene and dehydrated in ethanol. For Pgp, CYP46A1 and CH25H, antigen retrieval was not performed. For CYP27A1, slides were heated in citrate buffer via microwave for 10 minutes, followed by a cooling period of 20 minutes. Slides were blocked for endogenous peroxidase activity with 0.3% $H_2O_2$ in methanol for 10 minutes. Non-specific antigen binding was blocked with in Blocker$^{TM}$ Casein in TBS (37532, Thermo Fisher Scientific) for 30 minutes. Slides were incubated with antibodies for 1 hour. Antigen binding was visualised using Cell Signalling Technology secondary antibodies and DAB substrate kit. Nuclei were stained with Mayer's haematoxylin and washed with Scott's tap water. Sections were dehydrated with ethanol, washed with xylene and mounted onto coverslips with DePeX (Fluka; Gillingham, UK). The inter-observer intraclass correlation coefficient (ICC) was assessed between two independent scorers for each protein (AW, and one of two histopathologists, BW or LW) as as previously reported (Thorne *et al.*, 2018). The observers independently scored 10 matched triplicate tumour cores of the TMA (SF12B). The ICC found to range from 0.84 and 0.99, representing an 'almost perfect' to 'perfect' agreement (Barry *et al*, 2010). Scores recorded by AW were used for the analysis.

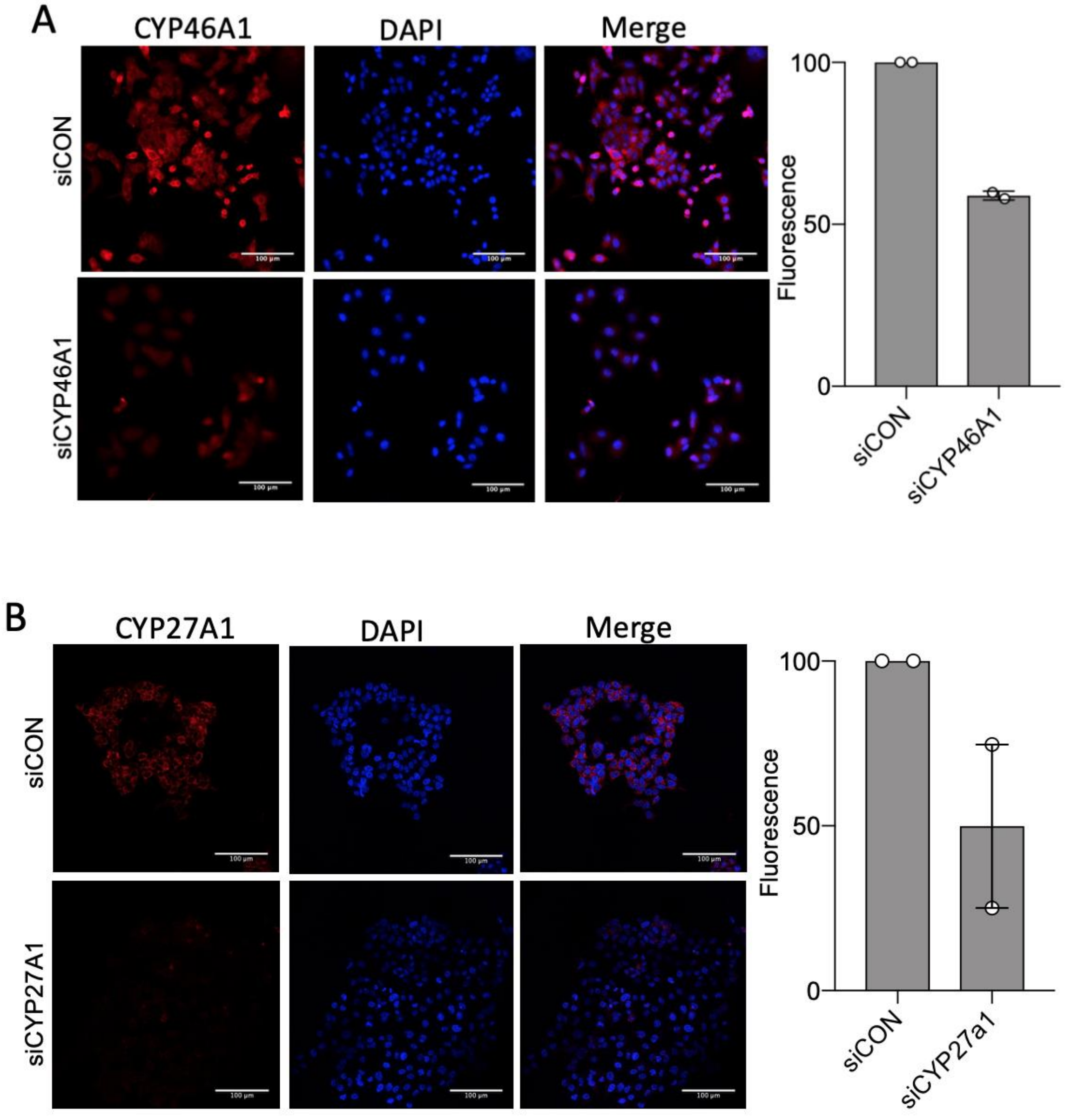


**Supplementary Figure S10. Validation of antibodies used for immunohistochemistry**

siRNA was performed in MCF7 cells against CYP46A1 (A) and in HEPG2 cells against CYP27A1 (B). Immunofluorescence signal was normalized to siControl (siCON) treated cells. Antibody signal is shown in red (left), DAPI nuclear stain shown in blue (middle) and merged (right). Scale bars show 100μM and images taken at x20 magnification. Fluorescence measured as mean gray value. Error bars represent SEM of 2 independent replicates each generated from at least three technical replicates.

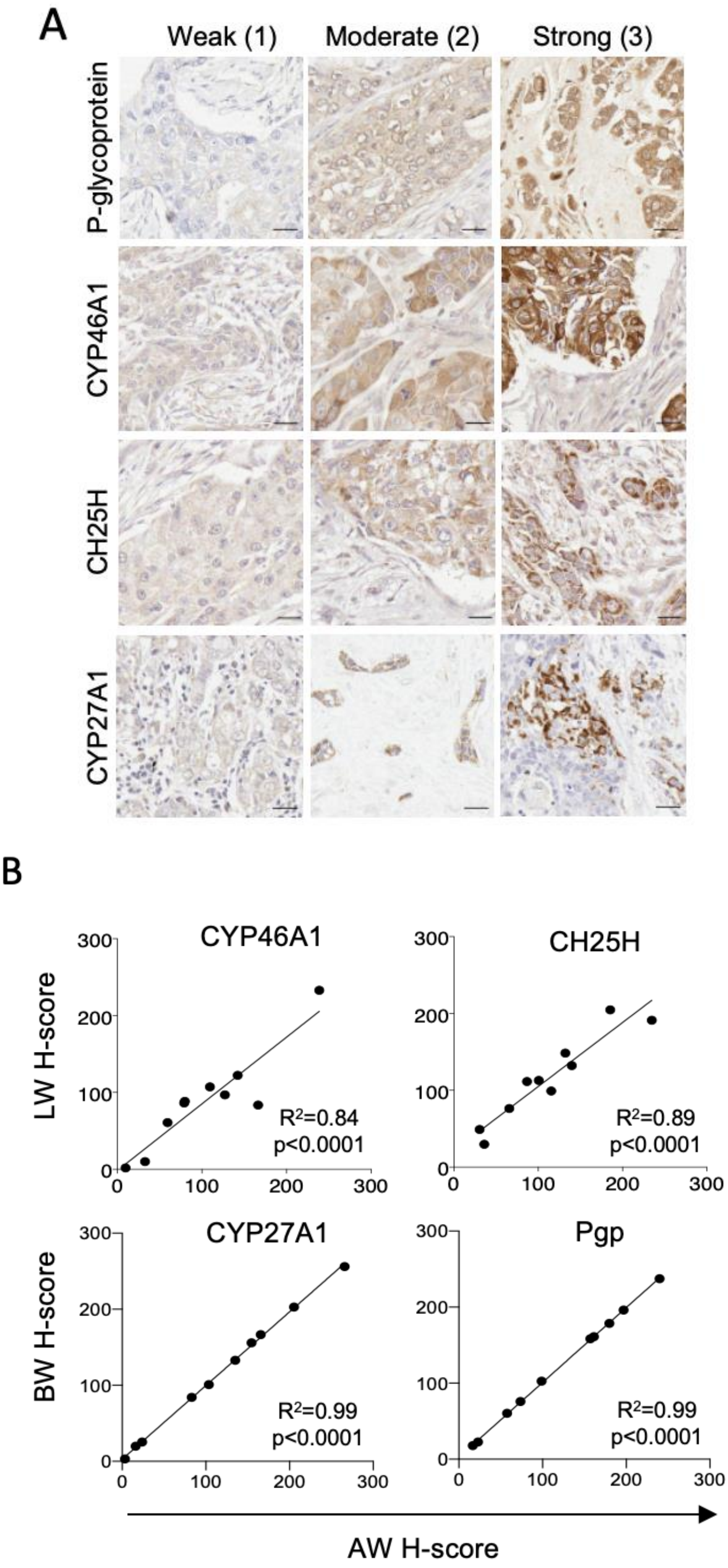


**Supplementary Figure S11: Immunohistochemistry scoring and validation for TMA**

Histoscore system for IHC showing examples of weak (1), moderate (2), and strong (3) staining for indicates proteins (A). Scale bars show 20μM and images taken at x40 magnification. Panel B shows interclass correlations between histoscores from two independent scorers for CYP46A1, CH25H, CYP27A1 and Pgp. Scores given by scorer 1, either BW or LW, are plotted against scorer 2, AW. Interclass correlation coefficients were generated through Pearson's correlations.

#### LC-MS/MS analysis of oxysterol concentration

Oxysterols were analysed as reported previously (Solheim *et al*, 2019) from 5mg homogenised tumour slices. Tumour slices were homogenized in 500 µl of internal standard solution (1.5 nm of deuterated 24OHC, 25OHC and 26OHC in 2-propanol) and 30 µl 6µM cholesterol-25,26,27-13C (Sigma-Aldrich) to monitor sample autoxidation using IKA T10 Ultra-Turrax homogenizer (VWR). 35 µl of 2M KOH (105032, Sigma-Aldrich, UK) in MeOH was added to 100 µl of homogenized sample solution and heated to 60° C for 120 minutes. Sterols were extracted through liquid-liquid extraction using 300 µl of n-hexane (VWR) and type 1 water mixture (1:1), retaining the light phase. Liquid-liquid extraction was performed a further two times, only adding 150 µl of n-hexane. Samples were evaporated in an Eppendorf concentrator plus and resuspended in 200 µl 2-propanol. Samples were eluted through an Oasis PRiME HLB 1 cc (30mg) SPE cartridge (186008055, Waters, US) with 200 µl MeOH, evaporated and then resuspended in 20 µl 2-propanol. Samples were incubated with cholesterol oxidase in 50 mM phosphate buffer pH7 for 60 minutes at 37° C. Samples derivatized using a 500 µl mixture of 15 mg Girard T reagent, 15 µl glacial acetic acid and 485 µl MeOH and left overnight at room temperature. Analyses were performed on a Dionex UltiMate 3000 UHPLC system connected to a TSQ Vantage triple quadrupole mass spectrometer. The instrument was equipped with an on-line HotSEP C18 SPE column to elute excess Girard P to waste. Separation was carried out by an AVE SuperPhenyl Hexyl xolumn (2.1 mm ID x 100 mm, core-shell) by a Dionex Ultimate 3000 UHPLC pump. Peaks were measured using Xcalibur (Thermo Fischer Scientific). To calculate concentration of metabolite per gram of tumour, a calibration curve was generated with 50–500pM standard mixtures containing 200pM internal standard solution. At least 10% of measurements were performed in duplicate by two independent researchers (SF13A). Representative chromatograms are shown in SF13B.

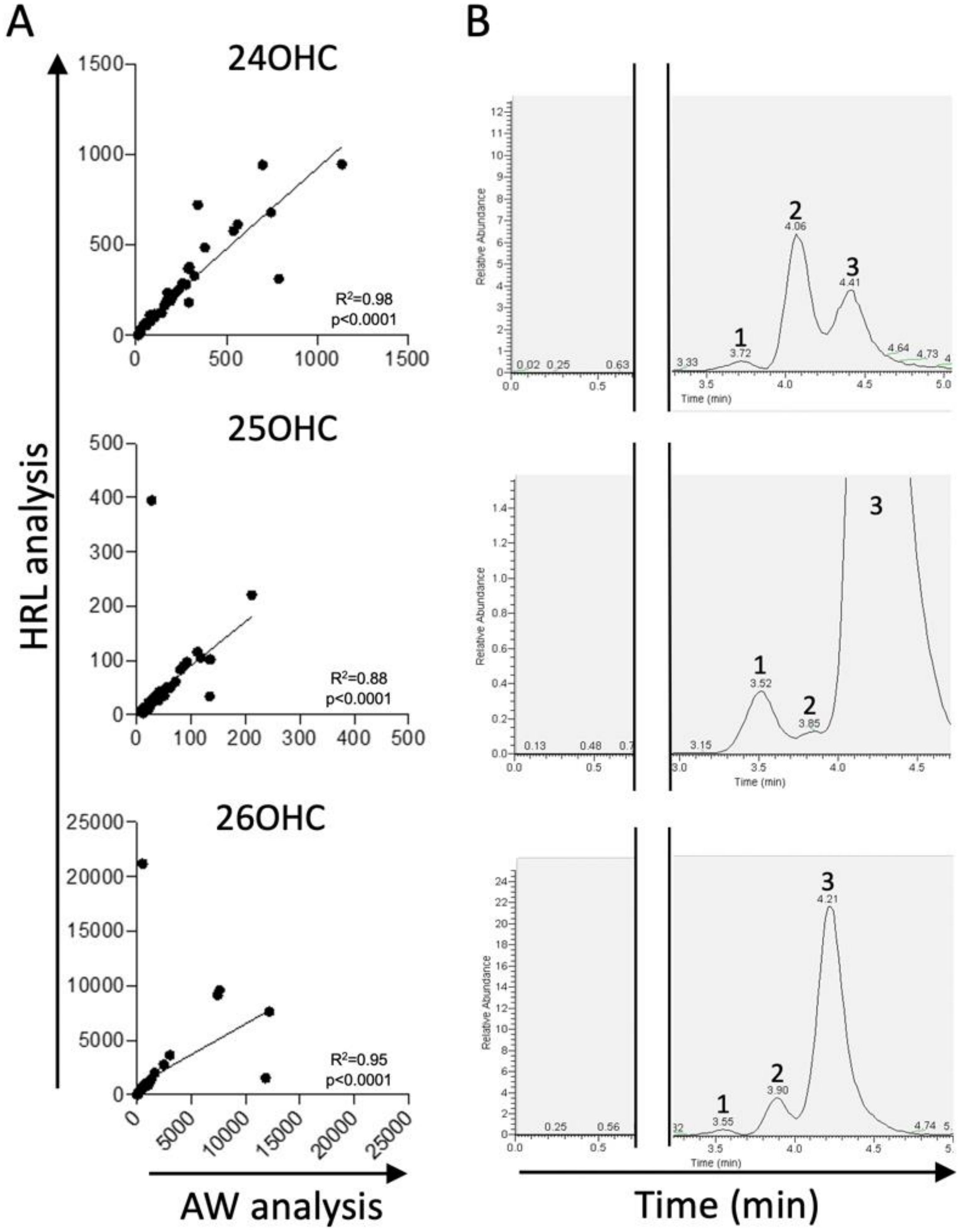


**Supplementary Figure S12: LC-MS/MS validation and example chromatograms**

Interclass correlations between two independent analyses measuring ion chromatograms (A). Interclass correlation coefficients were generated through Spearman's rank. Three example chromatograms (m/z 514.4 → 455.4) showing 25OHC (peak 1), 24OHC (peak 2), and 26OHC (peak 3).

#### Supplementary Table

| Characteristic | Category | Leeds Breast Research Tissue Bank No. of patients = 47 (%) | Tissue Microarray No. of patients = 146 (%) |
|---|---|---|---|
| Tumor Grade | 1 | 1 (2) | 2 (2) |
| | 2 | 9 (19) | 19 (13) |
| | 3 | 37 (79) | 123 (83) |
| | N/A | 0 (0) | 3 (2) |
| Tumour size | ≤ 35 mm | 27 (57) | 66 (45) |
| | > 35 mm | 20 (43) | 18 (12) |
| | N/A | 0 (0) | 64 (43) |
| Survival status | Alive | 25 (53) | 101 (68) |
| | Deceased | 22 (47) | 47 (32) |
| Recurrence status | None | 23 (49) | 102 (69) |
| | Present | 24 (51) | 46 (31) |

**Supplementary Table S1: Patient characteristics table**
